# A microRNA-based Johne’s disease diagnostic predictive system: preliminary results

**DOI:** 10.1101/2023.07.07.548088

**Authors:** Paul Capewell, Arianne Lowe, Spiridoula Athanasiadou, David Wilson, Eve Hanks, Robert Coultous, Michael Hutchings, Javier Palarea-Albaladejo

## Abstract

**Background:** Johne’s disease, caused by *Mycobacterium avium* subsp. *paratuberculosis* (MAP), is a chronic enteritis impacting welfare and productivity in cattle. Screening and animal removal are common for disease management, but efforts are hindered by low diagnostic sensitivity. Expression levels of small non-coding RNA molecules involved in gene regulation (microRNAs) altered during mycobacterial infection may present an alternative diagnostic method.

**Methods:** Levels of 24 microRNAs affected by mycobacterial infection were measured in sera from MAP-positive (n=66) and MAP-negative samples (n=65). They were used to train a collection of statistical and machine learning models to identify an optimal classifier for diagnosis.

**Results:** The best-performing model provided 72% accuracy, 78% AUC, 73% sensitivity and 71% specificity on average.

**Limitations:** Although control samples were collected from farms nominally MAP-free, low sensitivity in current diagnostics means animals may be misclassified.

**Conclusion:** MicroRNA profiling combined with advanced predictive modelling techniques accurately diagnosed Johne’s disease in cattle.

## Introduction

Johne’s disease is a chronic enteritis of ruminants caused by *Mycobacterium* avium subspecies *paratuberculosis* (MAP). It is of significant concern to the dairy industry as it can lead to decreased production, weight loss, and death in infected cattle (1). Estimates of losses range from £10m in the UK (2) to £158m in the US (3). Prevention and control of Johne’s disease is difficult as MAP is environmentally resilient and vaccine efficacy is variable and controversial (4). As such, management and control strategies focus on testing and targeted removal of infected animals (5). Despite detection being a central component of control, diagnosis can be challenging as clinical signs are insidious and non-specific (1). The most accurate diagnostic method to date is bacterial culture but is costly and can take months to yield results (6). The most commonly used tests are serum antibody ELISAs which have high specificity but show low sensitivity in cattle (≈0.44), especially in the early stages of infection, leading to low accuracy (≈0.55) (6,7). Furthermore, it has been shown that MAP shedding may be occurring for up to two years prior to positive milk ELISA testing resulting in high transmission rates that go undetected (8). Research is therefore needed to develop effective, early disease diagnostics and sustainable strategies to mitigate the economic impact of Johne’s disease.

A promising approach to develop novel and reliable diagnostics is to harness the information contained in the expression profiles of small non-coding RNA molecules. MicroRNAs (miRNAs) are found in most eukaryotes and play critical roles in gene regulation (9). They have diagnostic and prognostic value and are particularly suitable as biomarkers due to being stable and easily detectable in biofluids (10). Importantly, there have been several studies showing that expression levels of numerous miRNAs are altered by mycobacterial infection, including early-stage MAP (11). Diagnostic accuracy, or the capacity of a test to discriminate diseased and healthy samples, can be further enhanced through the deployment of advanced statistical and machine learning methods within an artificial intelligence (AI) framework. These analytical tools can significantly improve diagnosis by rapidly analysing experimental data, identify patterns that could be missed by humans or less sophisticated methods, and produce predictions to support an objective assessment of the uncertainty regarding infection status. They have already proven useful for improving diagnosis and prognosis in human medicine, for instance in oncology (12).

To explore the potential of miRNA profiling in Johne’s disease diagnosis, this preliminary study measured the expression of selected miRNAs impacted by mycobacterial infection in samples from MAP-infected and uninfected cattle. A curated collection of fifteen predictive models were thoroughly tuned, trained, and benchmarked to determine an optimal classifier for the diagnosis of Johne’s disease using miRNA profiles.

## Materials and Methods

### Sample Collection

All cases submitted to the UK-wide Premium cattle Health Scheme with reported signs of Johne’s disease between November 2021 and November 2022 were considered (SRUC Veterinary Services, Penicuik, UK). MAP-positivity was defined as animals with serum titers greater than 70% using an ID Screen Paratuberculosis Indirect ELISA Test (Innovative Diagnostics, Montpellier, France). In total, 66 MAP-positive cases were identified over the sampling period. For controls, 65 MAP-negative samples were randomly selected from the same scheme. To minimize false negatives, controls were selected from tier 1 risk herds with no Johne’s disease three years prior to sampling.

### MicroRNA Panel

MiRNAs to be included in the MAP-specific miRNA panel were selected through a review of 28 manuscripts identified in a PubMed-based search using the terms “(((Johne’s disease[Title/Abstract]) OR (Mycobacterium avium[Title/Abstract])) AND ((miRNA[Title/Abstract]) OR (microRNA[Title/Abstract])))”. The sequences of bovine miRNAs noted to have altered expression during MAP or mycobacterial infection were extracted from MIRBase (13). Signalling pathways targeted by each miRNA and the total panel were predicted using mirPathv3 and a p < 0.05 threshold (14). These sequences were used to design a custom panel for the Fireplex miRNA platform (Abcam, Cambridge, UK). Five miRNAs previously validated as standards in cattle were also included to act as normalizers for the expression data (Table 1).

**Table 1.**
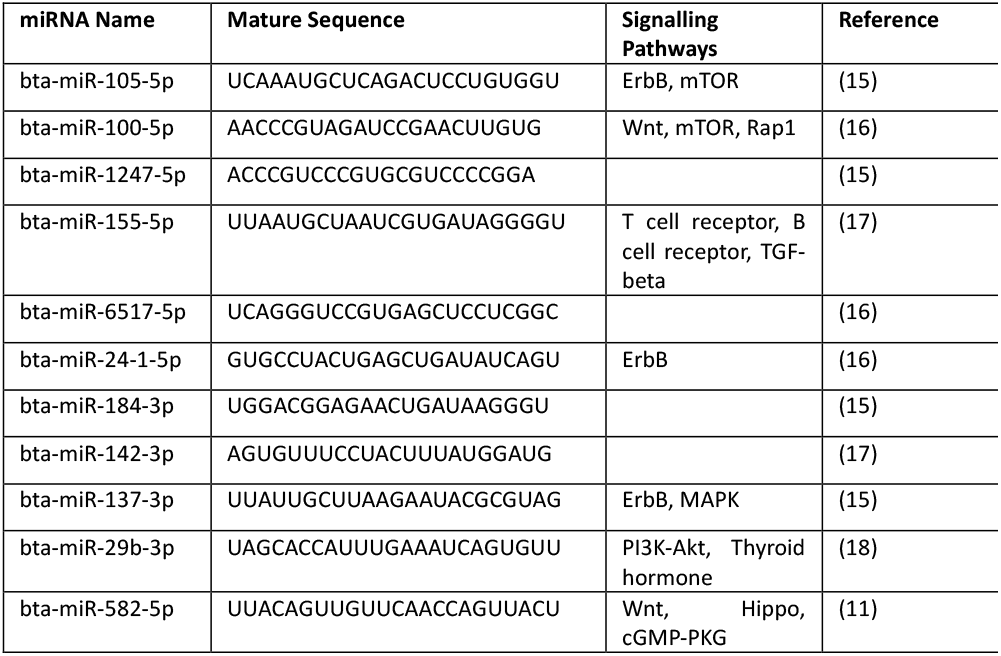
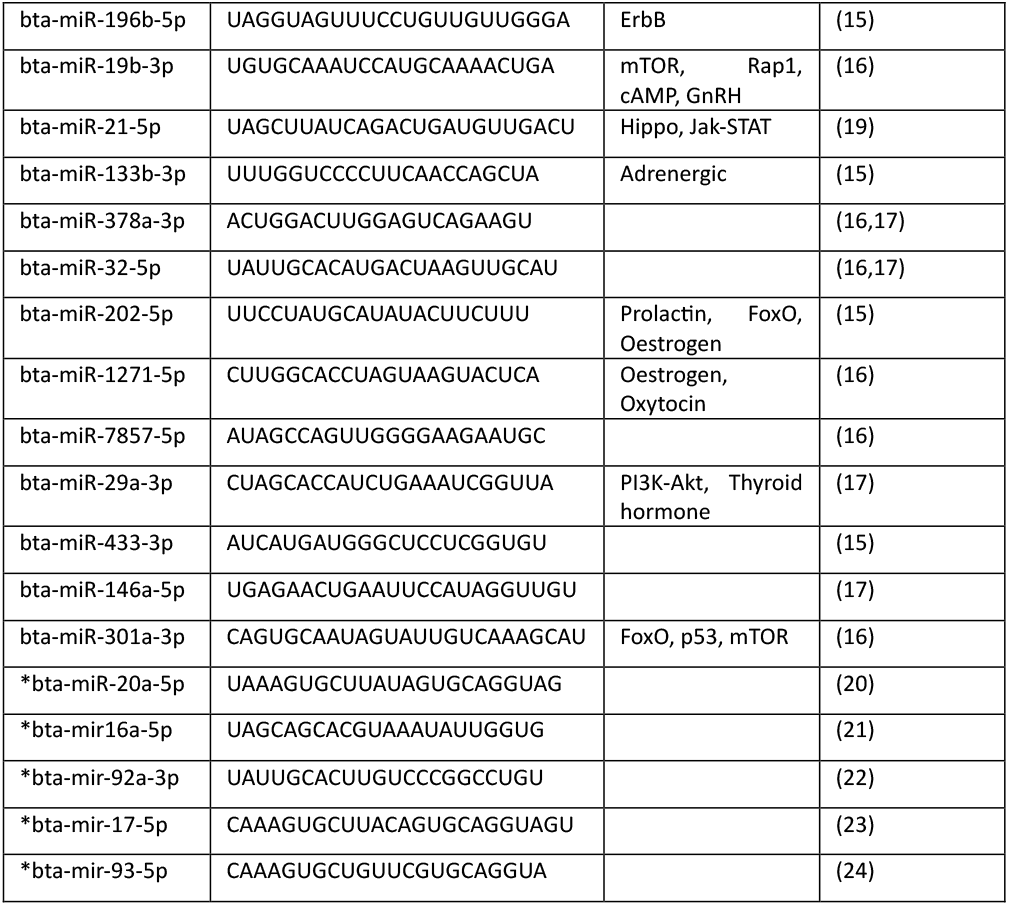
Summary information for the profiling panel indicating mature sequence, predicted signalling pathways targeted by each miRNA (p < 0.05). Normalizer miRNAs are indicated with an asterisk.

### Expression Analysis

To generate Fireplex miRNA profiles, 50 μL aliquots of sera from each sample were processed following the manufacturer’s instructions with hybridization, melt-off and capture temperatures of 39°C, 62°C and 39°C, respectively. Mean fluorescence intensities (MFI) of miRNA-specific particles per sample were measured using a Novocyte flow cytometer (Agilent, Santa Clara, CA, USA). Raw FCS files were exported to Fireplex Analysis Workbench 2.0.274 (Abcam, Cambridge, UK) and relative expression values prepared using the geNorm function with pre-selected normalizers.

### Predictive modelling

A collection of fifteen predictive classification models covering a range of popular linear, nonlinear and tree-based approaches were benchmarked: Bayesian generalised linear model (BayesGLM), boosted and penalised logistic regression (BootLogReg, PenLogReg), generalised partial least squares (GPLS), k-nearest neighbours (KNN), linear and quadratic discriminant analysis (LDA, QDA), neural network (NNET), ordinary, bagged and conditional inference recursive partition trees (RPART, TreeBAG, CPART), random forest (RF) and support vector machines (linear, SVM1; radial kernel-based, SVM2; radial and class weighting, SVM3). All models were trained on the Fireplex-processed and normalized (centred, scaled, Yeo-Johnson’s transformed) miRNA profiles as predictors and a binary response representing the diagnosis of Johne’s disease (infected vs. healthy outcome). Model training, including tuning of model parameters, was conducted through a 5-time repeated 10-fold cross-validation (CV) pipeline. The input data were randomly partitioned into ten folds, with nine folds used to train the model and one-fold used as validation set sequentially. Fold randomisation was repeated 5 times. This helped to balance any variance-bias trade-off in model fitting and, hence, contributed to a fairer assessment of their predictive ability with independent blind samples. Performance metrics included commonly applied methods, such as overall accuracy, area under the receiver operating curve (AUC), sensitivity and specificity. All measures ranged [0, 1], with values closer to 1 indicating better performance (95% confidence intervals given in parenthesis). Metrics were assessed with each validation set and then averaged, providing an assessment of how the model might perform on independent blind samples. This study was implemented on the R system for statistical computing v4.2.1 (25), using the specialised package caret v6.0-94. Extensive details about the models and methods used can be found in e.g. (26), (27) and (28).

## Results

A review of the literature suggested 24 miRNAs with altered expression in response to mycobacterial infection (Table 1), including experiments conducted with cattle and MAP (15,16,29). Across the total panel, there was enrichment for miRNA-target genes involved in PI3KT, oestrogen, ErbB and FoxO signalling (p < 0.05). After the MFI of each miRNA was measured across all samples, the mean coefficient of variation (CoV) was found to be 0.49 (0.38-60) for normalizing miRNAs and 1.23 (0.82-1.64) for diagnostically informative miRNAs. The lower CoV indicated that the miRNAs selected for normalization were stable and suitable standards. The cross-validated accuracies of the predictive models ranged in median between 0.57 (QDA) and 0.77 (RF) (Figure 1A). The best-performing model, RF, provided a mean accuracy of 0.72 (0.70-0.74), with this being statistically significantly greater than the naïve classifier accuracy (0.50; p < 0.001). Using infected status as reference class, the RF model provided an AUC of 0.78 (0.70-0.86) (Figure 1B), sensitivity of 0.73 (0.61-0.82), and specificity of 0.71 (0.59-0.80). Misclassification was more common for healthy samples (Figure 1C; 14.96% healthy classed as MAP-infected against 12.67% infected classed as healthy).

**Figure 1.**
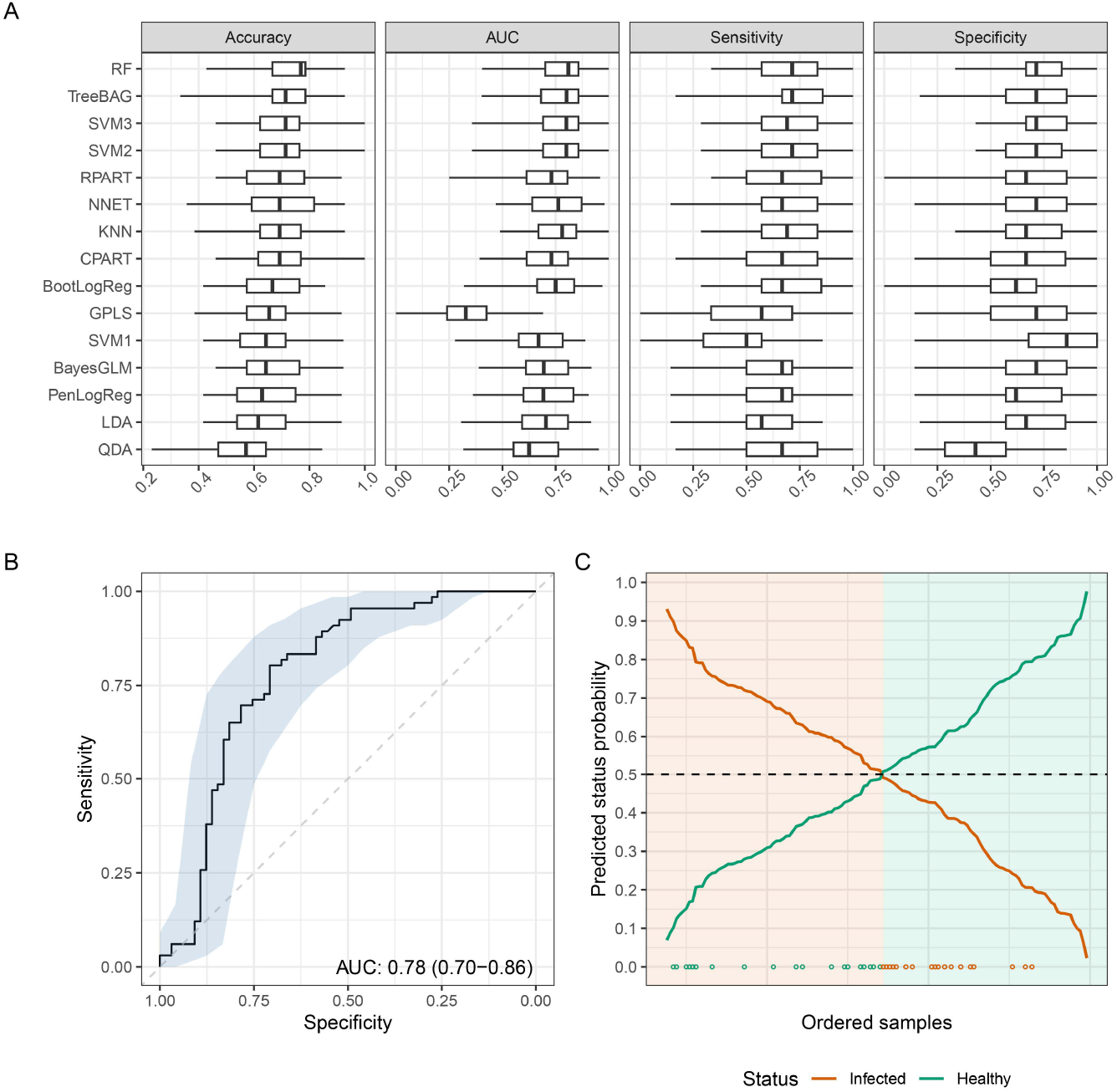
(A) Boxplots summarising common predictive performance metrics over 5-time repeated 10-fold cross-validation runs for the 15 trained models (ordered by overall classification accuracy from top to bottom). (B) Receiver operating curve (ROC) for the optimal random forest model and area under the curve (AUC). (C) Probabilities of status (infected/healthy) for collected cattle samples as predicted by the optimal random forest model (samples ordered from left to right according to probability). Sample classification based on the usual 0.5 probability threshold (dotted line), background colour indicates actual sample status. Points at the bottom indicate misclassified samples, with colour corresponding to their actual infectivity status.

## Discussion

The aim of this study was to evaluate the capacity for a miRNA expression panel to discriminate between MAP-positive and MAP-negative bovine sera.

The data indicate that a panel of 24 miRNAs, previously identified as having altered expression in MAP infection, was able to achieve an accuracy of 0.72 (sensitivity 0.73; specificity 0.71) in discriminating between healthy and MAP-infected cattle using the optimum Random Forest model within the provided sample population. It is possible that specificity (0.71) is under-estimated in our dataset as control animals were screened using an ELISA with low sensitivity (30). Efforts were taken to minimize this issue by only including control cattle from tier 1 risk farms but it was that noted misclassification was more common in controls (14.96% versus 12.67%). This may have been due to the presence of undiagnosed MAP-positive animals within these herds, given the low sensitivity of current ELISA screening methods. In future trials, cattle can be diagnosed using more accurate culturing techniques to overcome this shortcoming. Another possible issue may be diagnostic cross-reactivity with other mycobacterial species, such as *M. tuberculosis* (TB). Cross-reactivity can be negated by including MAP-or gut-infection specific miRNAs, but future studies will include challenging the model with known TB-positive sera. We hypothesise that comparing the miRNA profiles of MAP, TB and co-infected cattle will allow for the training of models able to discern species, as well as infection status.

In addition to widening the scope of the assay, accuracy may be improved by including additional informative markers. The current panel was suggested by a limited literature review and included target pathways known to be impacted by Johne’s disease, including PI3KT, oestrogen, ErbB and FoxO signalling (31,32). The panel could be expanded by including data from additional mycobacterial species, examining miRNAs known to target additional MAP-affected pathways or by performing a hypothesis-generating study using miRNASeq. MiRNAs may also improve the MAP diagnosis pipeline by exploiting their stability in biofluids, particularly milk (3,33). Most commercial tests for Johne’s disease use sera, adding expertise and cost barriers to screening. The feasibility of profiling milk miRNAs for MAP diagnosis is currently being evaluated. Finally, although primarily an infection of cattle, MAP also affects other ruminants (1). MicroRNAs are highly conserved between species, suggesting that the current miRNA panel would be suitable for sheep and goats with small adjustment, although species-specific trained models would be required. The capacity to provide a single diagnostic that allows for pathogen detection and identification across multiple species using an easily accessible biofluid makes miRNA profiling a highly attractive tool for veterinary diagnosis.

In summary, our preliminary study indicates that miRNA profiling in combination with advanced predictive modelling has the potential to serve as a diagnostic test for Johne’s disease in cattle. Efforts are currently ongoing to expand and validate the model with more sample data, which will further improve accuracy and expand the approach to characterise Johne’s disease stage, enhance early-stage diagnosis and distinguish between alternative mycobacterial pathogens through miRNA diagnostics methods. This technology is patent pending.

## Acknowledgements

This project was funded by Innovate UK (project number 10003360). EH and RC are employees of MI:RNA Ltd. PC and JPA are consultants for MI:RNA Ltd. SRUC acknowledges funding from the Scottish Government.

## Ethics

The use of historic sera was considered by the SRUC Ethical Committee and permission was granted as samples had been submitted with farmer’s consent via the Premium cattle Health Scheme.

